# Propagation and update of auditory perceptual priors through alpha and theta rhythms

**DOI:** 10.1101/2020.08.14.250514

**Authors:** Hao Tam Ho, David C. Burr, David Alais, Maria Concetta Morrone

## Abstract

To maintain a continuous and coherent percept over time, the brain makes use of past sensory information to anticipate forthcoming stimuli. We recently showed that auditory experience in the immediate past is propagated through ear-specific reverberations, manifested as rhythmic fluctuations of decision bias at alpha frequency. Here, we apply the same time-resolved behavioural method to investigate how perceptual performance changes over time under conditions of high stimulus expectation, and to examine the effect of unexpected events on behaviour. As in our previous study, participants were required to discriminate the ear-of-origin of a brief monaural pure tone embedded in uncorrelated dichotic white noise. We manipulated stimulus expectation by increasing the target probability in one ear to 80%. Consistent with our earlier findings, performance did not remain constant across trials, but varied rhythmically with delay from noise onset. Specifically, decision bias showed a similar oscillation at ~9 Hz that depended on ear congruency between successive targets. This suggests rhythmic communication of auditory perceptual history occurs early and is not readily influenced by top-down expectations. In addition, we report a novel observation specific to infrequent, unexpected stimuli that gave rise to oscillations in accuracy at ~7.6 Hz one trial after the target occurred in the non-anticipated ear. This new behavioural oscillation may reflect a mechanism for updating the sensory representation once a prediction error has been detected.

## Introduction

When observers make perceptual judgements on a sequence of images, their decisions are typically biased toward the immediately preceding stimulus. *Serial dependence*, as this sequential effect is known, has been demonstrated for a variety of stimuli, including orientation(Fischer & Whitney, 2014), numerosity (Cicchini et al., 2014), facial identity (Liberman et al., 2014) and gender (Taubert et al., 2016), and even beauty (Xia et al., 2016). Similar influences of stimulus history have been reported in audition for pitch discrimination (Arzounian et al., 2017; Chambers et al., 2017; Chambers & Pressnitzer, 2014). Although these biases can introduce perceptual errors, they are beneficial in the long run (Burr & Cicchini, 2014). As the world tends to remain constant over time, it can be advantageous to assume that the current sensory environment is similar to the one encountered just moments before. Serial dependence provides a fascinating window into the predictive mechanisms that the brain engages to overcome the noise and ambiguity inherent in sensory signals (Gregory, 1997; Von Helmholtz, 1910).

Many models of ‘predictive perception’ (Friston, 2005; Lee & Mumford, 2003; Rao & Ballard, 1999) assume that predictions, or *priors*, are generated in higher cortices and fed back to early sensory areas where incoming information is matched to the expectation. However, the mechanisms by which prior sensory information is communicated are still unknown. We (Ho et al., 2019) recently tested the hypothesis that recursive propagation and updating of stored prior experience are related to low-frequency neural oscillations (Arnal & Giraud, 2012; Engel et al., 2001; Friston et al., 2015). Using a novel, time-resolved behavioural method (Landau & Fries, 2012), we showed that auditory stimulus history biased perceptual decisions rhythmically through alpha oscillations at ~9 Hz (Ho et al., 2019). Here, we combine the same behavioural technique with an oddball paradigm (Fig. 1B) to investigate how sensory expectations are updated following a prediction error.

**Figure 1.**
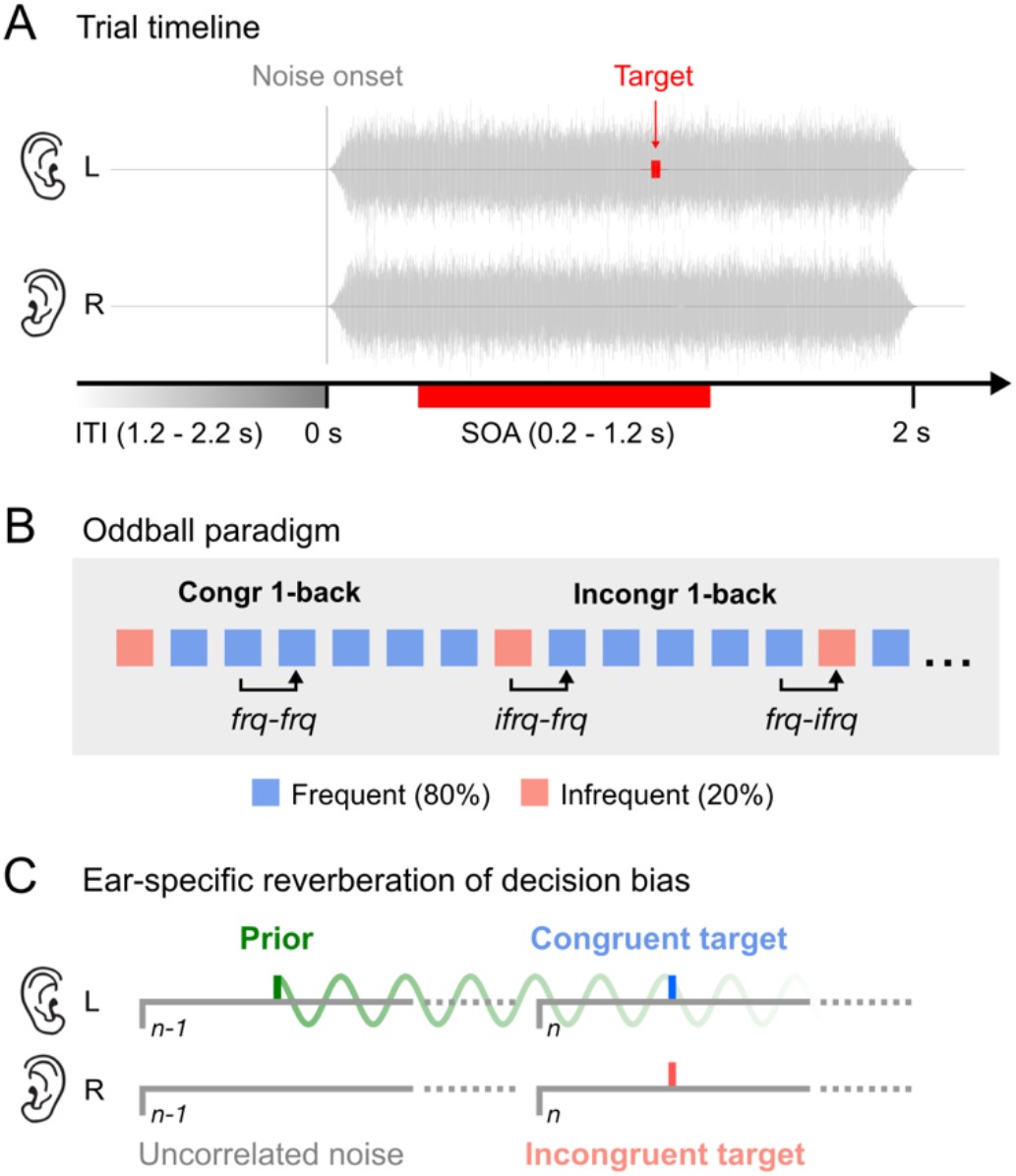
Experimental paradigm. **(A) Trial timeline.** Participants were required to identify the ear-of-origin of a brief (10 ms) monaural pure-tone (1 kHz) target embedded in dichotic, uncorrelated white noise that lasted for 2 s. The target occurred with a variable delay from noise onset, randomly drawn from a uniform distribution of stimulus-onset asynchronies (SOA) of 0.2-1.2 s. Each trial was followed by a silent intertrial interval (ITI) of 1.2-2.2 s. **(B) Oddball paradigm**. The experiment consisted of short blocks of 75 trials. We modulated attention by presenting the target to one ear for 80 % of the trials within each block. The order of L80 and R80 blocks (i.e., where the target occurred in the left and right ear for 80% of trials, respectively) were completely random. Participants were *not* informed about the manipulation at any point of the experiment. To test whether any rhythmic modulation of response bias was contingent on the ear congruency of successive targets, we separated the trials into three subgroups (i.e., *frq-frq, ifrq-frq*, *frq-ifrq*) depending on whether a frequent (*frq*) or infrequent target (*ifrq*) was presented on the previous (*t-1*) and current trial (*t*). **(C) Ear-specific reverberation of decision bias.** Oscillations of bias are observed when targets occur in the same ear (in this example, left ear) on consecutive trials (congruent, blue) but *not* when targets switch ears (incongruent, red). One possible explanation is that the previous target acts as a prior (green), biasing perceptual decisions on subsequent trials. Prior information is communicated rhythmically at alpha frequency.

Oscillations of perceptual performance (typically measured by accuracy) have been demonstrated with various visual tasks (Benedetto et al., 2016, 2018; Fiebelkorn et al., 2011; Fiebelkorn et al., 2013; Landau & Fries, 2012; Re et al., 2019; Tomassini et al., 2015). These rhythmic fluctuations are thought to arise from an attentional sampling mechanism involving slow-frequency brain rhythms in the theta (4-8 Hz) and alpha (8-12 Hz) frequency range (Fiebelkorn & Kastner, 2019; Landau, 2018; VanRullen, 2016). In an earlier study (Ho et al., 2017), we showed that auditory attention also samples information in a cyclic manner, switching between the two potential target locations (i.e., left and right ear) in a rhythmic fashion similar to vision (Fiebelkorn et al., 2013; Landau & Fries, 2012). Crucially, we used *signal detection theory* (Green & Swets, 1966; Macmillan & Creelman, 2004) to separate sensitivity and decision bias and found that in addition to the sensitivity oscillations at ~5-6 Hz, decision bias also oscillated but at significantly slightly higher frequencies, ~8-9 Hz. Furthermore, the oscillations in sensitivity exhibited an antiphase relationship between the ears while bias oscillations did not, suggesting that rhythmic fluctuations of sensitivity and decision bias over time arise from separate mechanisms.

A number of EEG studies in vision have reported modulations of decision bias by alpha rhythm (Iemi et al., 2017; Limbach & Corballis, 2016; Samaha et al., 2017) linked to stimulus expectation (Sherman et al., 2016) and perceptual choice history (Lange et al., 2013). Our finding in auditory behaviour that past perceptual information is propagated via alpha oscillations (Ho et al., 2019) is consistent with these results. Specifically, we showed that rhythmic modulations of decision bias at ~9.4 Hz were contingent on the previous target occurring in the same ear as the current one. At the same time, we observed a long-lasting serial dependence in decision bias that was clearly linked to the oscillation. These fluctuations in bias could be related to the “perceptual echo”, a long-lasting (up to 1 s) reverberatory response shown to be involved in short-term memory processes (Chang et al., 2017; VanRullen & Macdonald, 2012). Interestingly, Lozano-Soldevilla and VanRullen (2019) recently presented evidence that perceptual echoes are *travelling waves,* behaving like “radar beams” that “scan” areas of the brain, from posterior to frontal, in cycles of ~100 ms. The perceptual system may exploit these characteristics to exchange information across brain regions.

Several studies have tried but failed to find evidence for similar echo responses in audition (İlhan & VanRullen, 2012; Zoefel & Heil, 2013). However, these null findings do not necessarily mean that the auditory system samples sensory information differently than vision (VanRullen et al., 2014) and could instead be attributed to the general difficulty of recording alpha oscillations in audition with non-invasive EEG (Weisz et al., 2011). In support of auditory alpha oscillations, intracranial recordings (Lehtelä et al., 1997; Sedley et al., 2015) point unequivocally to the existence of auditory alpha with similar properties as in vision (Weisz et al., 2011), and our own studies (Ho et al., 2017, 2019) show that alpha oscillations can be readily demonstrated in audition behaviourally.

In the current study, we applied the same time-resolved behavioural method (Ho et al., 2017, 2019) to examine how alpha oscillations are engaged in predictive processes under high stimulus expectation. To keep the current experiment comparable to our previous study (Ho et al., 2019), we employed the same task, which required participants to identify the ear-of-origin of a brief monaural pure-tone target masked by uncorrelated dichotic white noise. To bias listeners’ expectation (and attention) toward one ear, we used an oddball paradigm where the target occurred with 80 % probability in one ear and 20 % in the other. Importantly, this paradigm also allowed us to examine the effect of unexpected stimulus events on auditory perceptual performance over time.

Abrupt perceptual changes rarely occur in the real world. Nevertheless, the sensory system must be prepared to revise its predictions in the face of an error. A great amount of EEG research has been devoted to understanding the neural mechanisms underlying regularity formation and violations thereof. One electrophysiological response that is typically evoked by infrequent, unexpected auditory and visual events is the Mismatch Negativity (MMN) (Näätänen et al., 2001, 2007). Extensive research on this event-related potential (ERP) response suggests that the MMN is elicited by a mechanism responsible for detecting violations of perceptual regularities and updating sensory predictions (Garrido et al., 2009; Stefanics et al., 2014; Wacongne et al., 2012; Winkler, 2007; Winkler et al., 2009). More recently, a number of findings suggest that generation of the MMN depends on oscillatory activity in the theta frequency band (Choi et al., 2013; Fuentemilla et al., 2008; Garrido et al., 2008; Hsiao et al., 2009; Ko et al., 2012). Here, we explore the possibility of an oscillatory correlate of the MMN in behaviour.

## Materials and Methods

### Participants

Seventeen healthy adults (5 male and 2 left-handed) with normal hearing took part in the experiment. The mean age was 24 ± 7 years. All participants provided written, informed consent. The study was approved by the Human Research Ethics Committees of the University of Sydney.

### Experimental procedure

Participants sat in a dark room and listened to auditory stimuli via in-ear tube-phones (ER-2, Etymotic Research, Elk Grove, Illinois) with earmuffs (3M Peltor 30 dBA) to isolate external noise. They kept their eyes open, and the computer screen was completely black throughout the testing. On each trial, 2 seconds of binaural broadband white-noise (randomly generated each trial and uncorrelated between the two ears) were presented together with the monaural target tone. The noise burst served to reset potential oscillations, similar to a visual or auditory cue (Fiebelkorn et al., 2011; Fiebelkorn et al., 2013; Landau & Fries, 2012; Re et al., 2019) and action or saccadic execution (Benedetto et al., 2016, 2018; Benedetto & Morrone, 2017; Tomassini et al., 2015, 2017). During the 2-s noise burst, within 0.2–1.2 s from noise onset, the target (1000 Hz, 10 ms) was delivered randomly with a probability of 80 % to one ear and 20 % to the other ear. For each ear, the target intensity was kept near individual thresholds (75 % accuracy), using an accelerated stochastic approximation staircase procedure (Faes et al., 2007; García-Pérez, 2011). Participants reported the *ear of origin* of the tone via button press (ResponsePixx, Vpixx Technologies, Saint-Bruno, Quebec). The next trial started after a silent inter-trial interval (ITI) of random duration ranging from 1.2–2.2 s. Participants completed 4,200 trials in total, divided into 60 blocks of 70 trials each. On half of the blocks, the target occurred with 80 % probability in the left ear (henceforth, *L80*) and for the remainder of the blocks, 80 % in the right ear (*R80*). Participants were not informed about the experimental manipulation at any stage of the experiment. They completed a practice block of 20 trials with feedback at the start of the experiment, but no feedback was given during the actual testing. Stimuli were presented using the software *PsychToolbox* (Brainard, 1997) in conjunction with *DataPixx* (Vpixx Technologies) in MATLAB (Mathworks, Natick, Massachusetts). Trials were excluded if the response occurred before the target onset or after the noise offset, or if the reaction time (RT) exceeded the 99 % confidence interval of that individuals’ RT distribution (fitted with a gamma function).

### General linear model (GLM)

To examine the individual data for oscillations and evaluate their coherence across subjects, we used an approach based on single trials (Benedetto et al., 2018; Tomassini et al., 2017). We have applied the same method in our previous study (Ho et al., 2019). The response *y*_i_(*i* = 1, 2… *n*, where *n* is the total number of trials) to a target at time *t*_*i*_ (i.e., the interval from noise onset to target onset in seconds) is modelled by the linear combination of harmonics at each tested frequency (*f*) as follows:

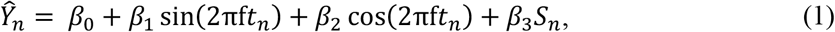

where 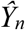 represents the predicted responses, *S*_*n*_ is the stimulus, which takes the value –1 or +1 for left and right target, respectively, and *β*_0_, *β*_1_, *β*_2_ and *β*_3_ are fixed-effect regression parameters, estimated using the *linear* least-squares method of MATLAB (*fitlm* function from the *Statistics and Machine Learning* toolbox). The fit was performed for each frequency *f* in the range from 4 to 20 Hz with 0.1Hz steps. Each trial’s accuracy and response were coded as follows: *y*_i_ = 1 for correct and *y*_i_ = −1 for incorrect responses, and for response bias, *y*_*i*_ = 1 for a ‘right’ response and −1 for ‘left’. As non-linear transformation of 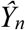 made no difference in our study (for further discussion, see Knoblauch & Maloney, 2008), we did not model our binary responses with a logit or probit link function. With linear transformation and without the *S*_*n*_ regressor the method is mathematically equivalent to fitting the discrete binary data with a sinewave function. Hence, it provides a good approximation of the Fourier spectra of the time series.

### Statistical methods

The significance of the model fit in Eq. 1 was evaluated using a two-dimensional (2D) permutation test: we shuffled the SOAs of each individual’s trials to create 1,000 surrogate datasets per subject and fitted each dataset with the same model described in Eq. 1. As with the original data, the resulting *β*_1_ and *β*_2_ were averaged across subjects for every frequency tested. This yielded a joined distribution of 1,000 surrogate means for each frequency from 4-12 Hz in 0.1-Hz steps. To correct for *multiple comparisons*, we determined the *maximal vector* across all tested frequencies for every permutation. This resulted in a joint distribution of 1,000 maximal vectors, against which we compared the original group mean.

In cases where we wanted to test a specific frequency *post-hoc*, we applied the one-sample and paired-samples Hotelling’s *T*^2^ test which are multivariate equivalents of the one-sample and paired-samples *t*-tests (Benedetto et al., 2018; Tomassini et al., 2017):

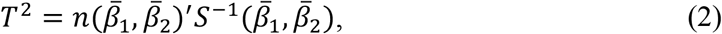

where *n* denotes the sample size and *S*^−1^ is the inverse of the covariance matrix. In the one-sample test, 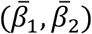 is the sample mean and in the paired-samples test 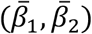 the mean sample difference. For small samples (i.e., *n* < 50), Hotelling’s *T*^2^ is transformed as follows:

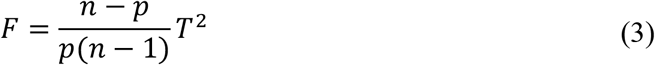

with *p* and *n-p* degrees of freedom (Hotelling, 1951).

The error in amplitude and phase across participants was estimated by computing the average 2D scatter of the individual data points using:

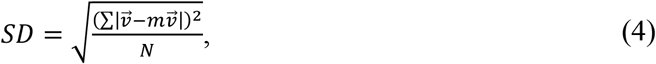

where *N* is the number of individual vectors. We obtained the amplitude SEM by projecting the SD values along the average vector 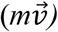 and divided them by the 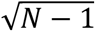. A similar procedure was applied to calculate the phase SEM, using propagation of errors.

## Results

### Rhythmic modulations of response bias but not of accuracy

Participants identified the ear-of-origin of a brief monaural target embedded within dichotic white noise (Fig. 1A). To bias expectation toward one ear, we presented the target with 80 % probability to one ear and 20 % to the other (Fig. 1B). First, we tested for rhythmic modulations of response bias and accuracy by low-frequency oscillations within the theta (~4-8 Hz) or alpha range (~8-12 Hz) for all data, irrespective of whether the target was presented more frequently to the left or right ear. For this analysis, we used the same general linear model (GLM) as in our previous studies (Benedetto et al., 2018; Ho et al., 2019; Tomassini et al., 2017), as this method yields equivalent results to the *curve-fitting* method that can also be applied to test for rhythmic fluctuations in behavioural data (Benedetto et al., 2016; Ho et al., 2017). The advantage of the GLM method is that it does not require grouping of individual responses into arbitrarily defined time bins but fits a linear model (Eq. 2 in Methods) to the individual responses to estimate the sine and cosine parameters for each participant and tested frequency (see Methods for more details). The results of the GLM are summarised in Figure 2.

**Figure 2.**
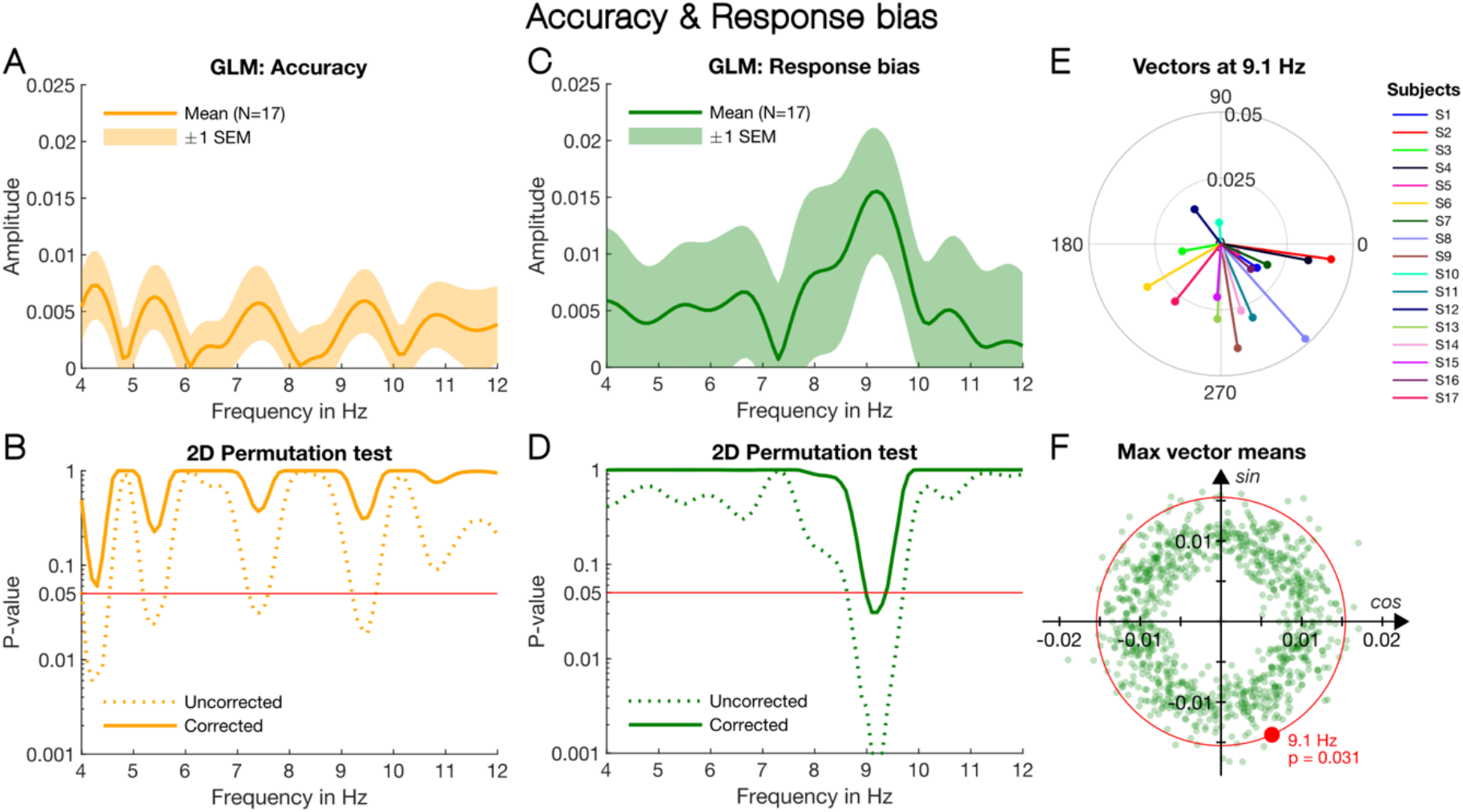
GLM results for response bias and accuracy. **(A)** Amplitude spectrum of accuracy: The amplitudes were computed with the beta coefficients obtained from the GLM which was fitted to the responses collapsed across the two target-probability conditions, L80 and R80. The orange line shows the mean vector length of the individual subject vectors (N=17) at every frequency from 4 to 12 in 0.1-Hz steps. The orange shaded area around the line indicates the dispersion of the individual subject vectors for the same frequencies and shows ± 1 standard error of the mean (SEM). **(B)** Results of 2D permutation test for accuracy: The orange solid line shows the p-values yielded by the 2D permutation test, corrected for multiple comparisons (see Methods). The uncorrected *p-*values are shown by the orange dotted line. The red line indicates *α* = 0.05. No frequency survived the strict multiple comparison correction we applied. **(C)** Amplitude spectrum of response bias: As in (A), we collapsed responses across the L80 and R80 conditions. The green line shows the mean vector length of the individual response bias vectors with the shaded area around indicating the vectors’ dispersion in ± 1 SEM. There is a one clear peak in the amplitude spectrum around 9.1 Hz. **(D)** Results of the 2D permutation test for response bias. Only the frequencies around 9.1 Hz remain significant after the correction. **(E)** Response bias vectors at 9.1 Hz for individual subjects. **(F)** Illustration of the 2D permutation test, corrected for multiple comparisons. Each green dot represents the maximal mean vector across all tested frequencies (frequency changes for the individual dots), obtained by permuting the individual response data. The red dot indicates the maximal mean vector of the original group data at 9.1 Hz. The *p*-value reflects the proportion of permuted vectors equal or exceeding the original mean vector (i.e., all the green dots that fall outside the red circle).

The amplitude spectra for accuracy and response bias in Figure 2A&C were obtained by computing the vector mean (orange and green lines) of the individual subject vectors (plotted in Fig. 2E for response bias at 9.1 Hz) for each tested frequency (4-12 Hz in 0.1-Hz steps). The shaded areas around the lines indicate ± 1 SEM. The bias spectrum in Fig. 2C shows a clear peak around 9.1 Hz with an amplitude of *A* = 0.015 ± 0.006. Using a permutation test (see Fig. 2F), shuffling the individual response 1,000 times and fitting the same linear model to the surrogate data, we confirmed that the peak at 9.1 Hz was statistically significant, *p* = 0.03. As shown in Figure 2D, at no other frequency did the *p*-values corrected for multiple comparisons (green solid line) approach significance (red line). Figure 2F illustrates how we computed the corrected *p*-values. Each green dot represents the maximal mean vector *across all tested frequencies*, determined after each shuffling. The red dot represents the maximal mean vector length of the original individual data (i.e., the mean vector at 9.1 Hz) computed by averaging the individual subject vectors shown in Fig. 2E. The *p*-value reflects the proportion of maximal permutation vectors that exceeds the original data. In practice, this means all the green dots that fall outside the red circle in Fig. 2F. Note in Fig. 2E how coherently the individual subjects cluster around a mean phase angle of *φ* = 294° ± 9° SEM.

The amplitude spectrum for accuracy is shown in Fig. 2A. In contrast to response bias (Fig. 2C), there are several large peaks in the accuracy spectrum. However, none of the peaks survived the significance test with multiple comparison correction, as shown in Figure 2B by the orange solid line corresponding to the corrected *p*-values. The absence of a significant oscillation frequency for accuracy is consistent with our previous study (Ho et al., 2019) and is possibly due to the antiphase relationship between the left- and right-ear sensitivity observed in our earlier study (Ho et al., 2017). We tested whether the accuracy data in the L80 and R80 conditions have different phases in a post-hoc analysis reported below (Fig. 3D).

**Figure 3.**
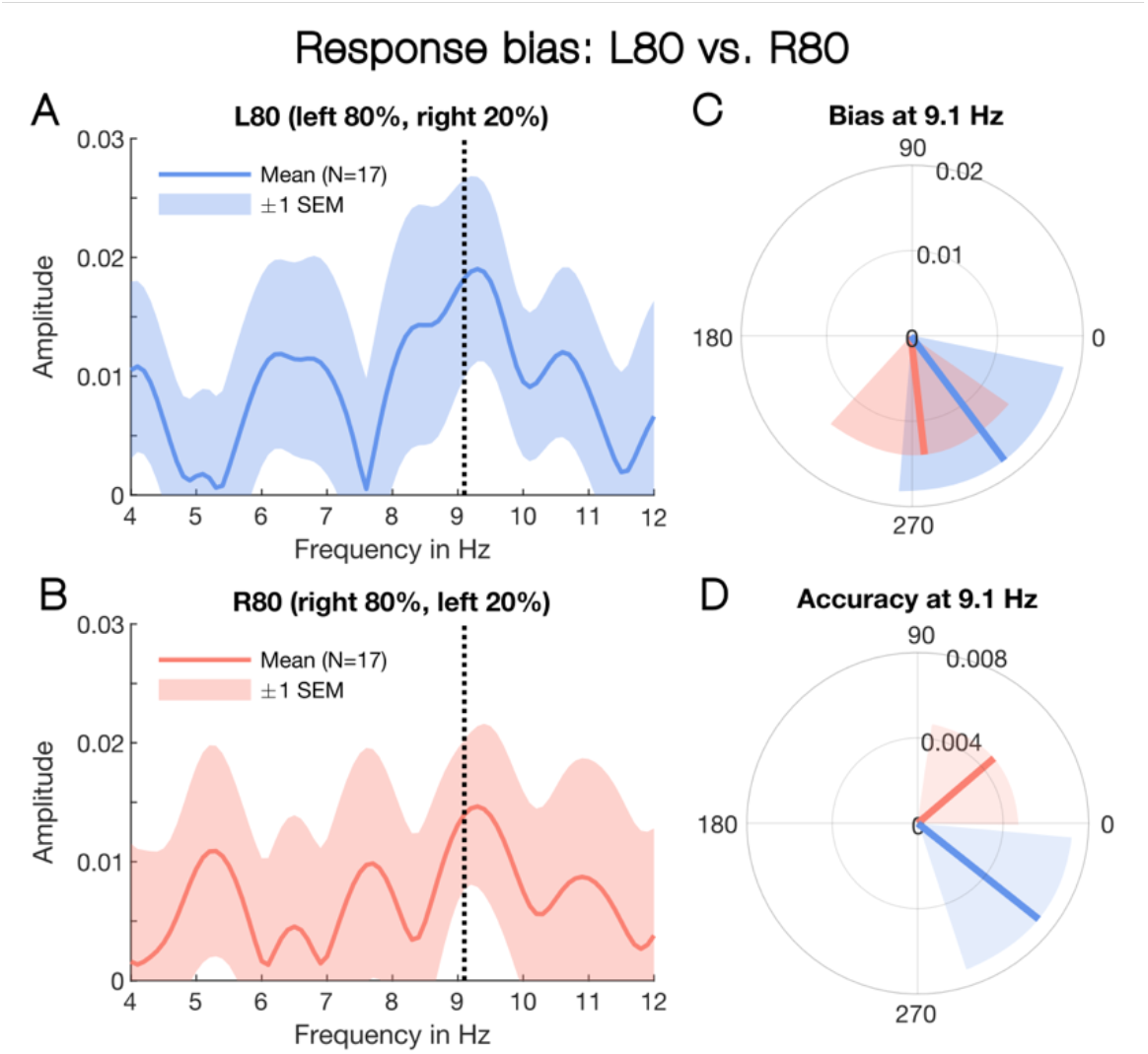
Response bias separated for L80 and R80. **(A)** Amplitude spectrum for the L80 condition: The blue line represents the mean vector length across subjects (N=17) at each tested frequency. The shaded area around the blue line reflects the dispersion of individual vectors measured in ± 1 SEM. The dotted line indicates 9.1 Hz, the peak frequency identified in Fig. 2C. **(B)** Amplitude spectrum for the Right-80 condition. As in (A), the shaded area surrounding the red line represents ± 1 SEM, and the dotted line indicates the peak frequency, 9.1 Hz. **(C)** The vector means for at 9.1 Hz in the L80 (blue line) and R80 conditions (red line) for response bias. The shaded areas in blue and red indicate ± 1 SEM and clearly overlap. **(D)** The vector means for accuracy at 9.1 Hz. As in (C), the red and blue lines represent the L80 and R80 conditions. Note, in contrast to bias, the shaded areas representing ± 1 SEM do not overlap.

### Similar alpha oscillations of bias whether attention sustained at left or right ear

Having shown that response bias fluctuates at alpha rhythm (Fig. 2C&D), we verified *post-hoc* that this 9.1-Hz oscillation was present in both target-probability conditions, i.e., L80 and R80. To do so, we separated the responses according to the two conditions and applied the same GLM method to each dataset, testing specifically the peak frequency, 9.1 Hz (Figs. 2C-F). Figure 3A shows the amplitude spectrum for response bias in the L80 condition. The amplitude reaches maximum at 9.3 Hz. At 9.1 Hz, the amplitude measures 0.018 ± 0.008, with a mean phase of *φ* = 306° ± 10°. In the R80 condition (Fig. 3B), the amplitude peaks also at 9.3 Hz. At 9.1 Hz, *A* = 0.014 ± 0.006 and *φ* = 276° ± 11° SEM. Using the paired-sample Hotelling *T*^2^ test, a multivariate equivalent of the univariate *t*-test, we verified that the amplitude of the response bias oscillation in the L80 and R80 conditions was not significantly different at 9.1 Hz, *T*^*2*^ = 3.30, *F*_*2, 31*_ = 1.23, *p* = 0.32 (Fig. 3C). In addition, we applied the one-sample Hotelling *T*^*2*^ test and confirmed that the response bias amplitude in both the L80 and R80 was significantly different from zero at 9.1 Hz, *T*^*2*^ = 10.95, *F*_*2,15*_ = 5.13, *p* = 0.02 and *T*^*2*^ = 11.08, *F*_*2, 15*_ = 5.19, *p* = 0.019, respectively. Taken together, the results suggest that bias fluctuated at the same alpha frequency and phase, irrespective of whether the target occurred more often in the left or right ear.

For comparison, we examined the individual accuracy vectors in the L80 and R80 vectors at 9.1 Hz and found that the vectors in the two conditions pulled away from each other, with L80 showing a mean phase angle of 322° ± 8° and R80 a mean phase angle of 40° ± 10° (Fig. 3D). Again, using the paired-sample Hotelling *T*^*2*^ test, we tested the difference between the L80 and R80 accuracy at 9.1 Hz. The result approached significance with *T*^*2*^ = 7.66, *F*_*2, 31*_ = 3.59, *p* = 0.053.

### Rhythmic fluctuation of bias is contingent on the previous stimulus

Our previous study (Ho et al., 2019) showed that rhythmic fluctuations in response bias were contingent on the target of the previous trial occurring in the same ear as the current target (illustrated in Fig. 1C). To test whether the oscillation of bias observed in the present study was similarly contingent on ear congruency between successive targets, we ran a one-back analysis on the response data. As illustrated in Fig. 1B, we divided the trials into three subsets, depending on whether the stimulus on the current trial (*t*) and previous trial (*t-1*) was a frequent (*frq*) or infrequent target (*ifrq*). Because infrequent targets very rarely occurred successively, we did not analyse those trials but focused on the remaining three conditions: (i) frequent preceding frequent (*frq-frq*), (ii) infrequent preceding frequent (*ifrq-frq*) and (iii) frequent preceding infrequent (*frq-ifrq*). We expected that response bias should oscillate at around 9 Hz in the *frq-frq* data where the trials contained *congruent* targets but not in the *frq-ifrq* data where the targets switched ears on consecutive trials. Because further findings from our previous (Ho et al., 2019) work suggested that reverberations of response bias last at least two trials, we anticipated a 9-Hz oscillation in the *ifrq-frq* data, as these trials are most likely preceded by a frequent target two trials back.

Figures 4A&C show the response bias amplitude spectra for *frq-frq* and *ifrq-frq* trials. Both spectra contain a clear peak near 9 Hz. For the *frq-frq* data (Fig. 4A), the amplitude is maximal at around 9.4 Hz, *A* = 0.015 ± 0.007 SEM, and for the *ifrq-frq* data around 8.8 Hz, *A* = 0.03 ± 0.017 SEM. We show the whole spectrum (i.e., 4-12 Hz) for completeness, however, our hypothesis that oscillations of response bias depend on ear congruency was specific to frequencies near 9 Hz. Figure 4G summarises the statistical results obtained by the same 2D permutation test as in Fig. 2B&D (dotted lines, not corrected for multiple comparisons). The red line indicates *α* = 0.05. There is one significant frequency common to both 1-back conditions, *frq-frq* (dark-green solid line) and *ifrq-frq* (green dotted line), and that is 9.2 Hz, *p* = 0.041 and *p* = 0.044, respectively. The individual vectors for *frq-frq* and *ifrq-frq* at 9.2 Hz are plotted in Figure 4B&D. Although, the vector dispersions in the *frq-frq* and *ifrq-frq* conditions are greater than in Fig. 2E, the vectors cluster around a mean phase close to that in Fig. 2E, with *frq-frq* showing a mean direction of 264° ± 11° and *ifrq-frq* 283° ± 13°. To verify that the oscillations in the *frq-frq* and *ifrq-frq* data reflect the same oscillation, we pooled together the trials from the two conditions and ran the same GLM analysis on the resulting dataset. Bias oscillated clearly at ~9.2 Hz, *p* = 0.004 (uncorrected). The amplitude was *A* = 0.015 ± 0.006 SEM and the phase 272° ± 10°.

**Figure 4.**
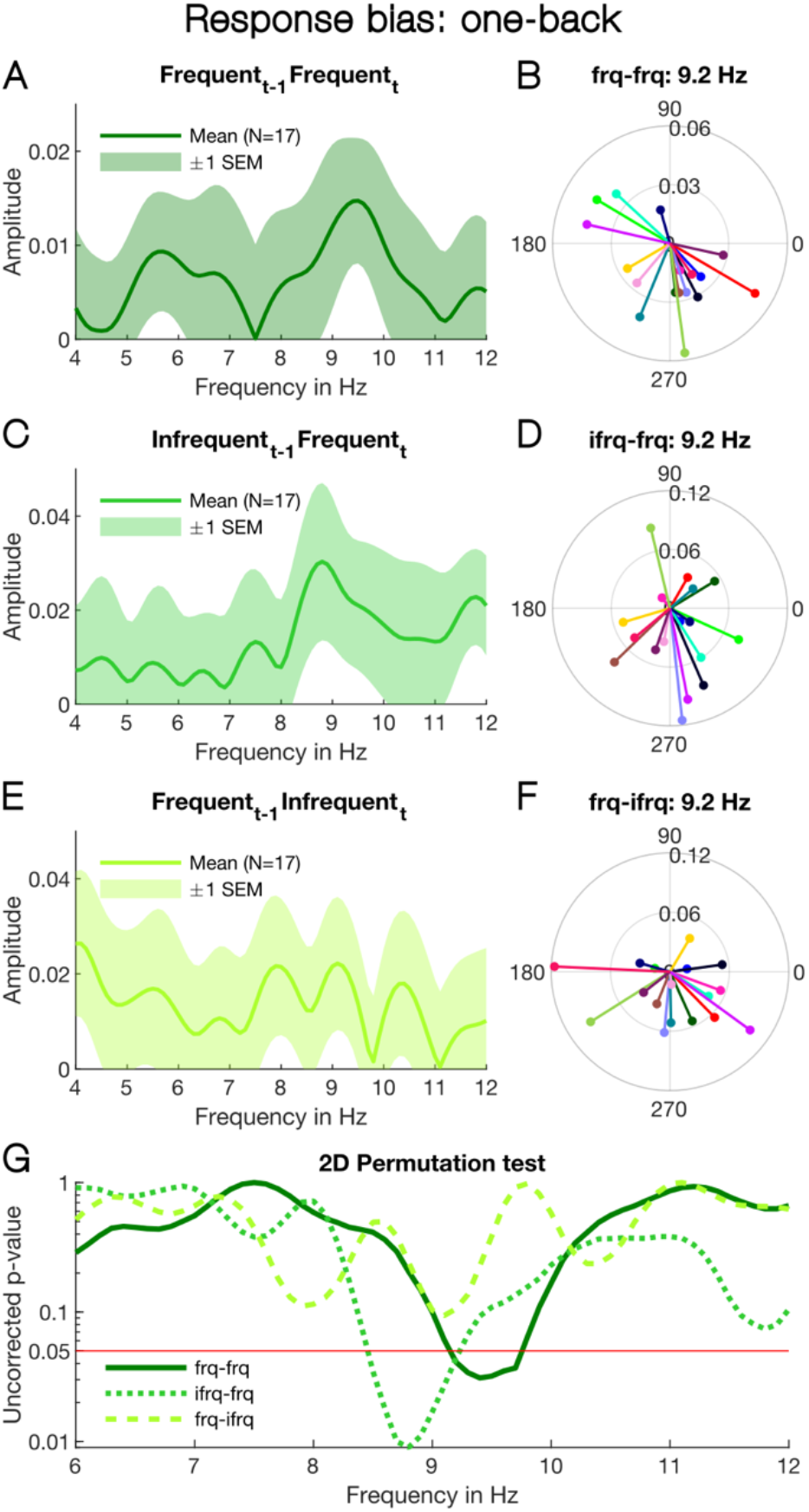
One-back analysis of response bias. **(A)** Amplitude spectrum of response bias for trials where frequent targets occur on consecutive trials: The dark-green line shows the mean vector length across individual subjects at the tested frequencies and the shaded area indicates ± 1 SEM (N =17). Although in this analysis we are only interested in the frequencies around 9 Hz, we show the whole spectrum, i.e., 4-12 Hz, for completeness. The mean number of trials in the *frq-frq* condition was 1141 ± 78. **(B)** Vectors for individual subjects at 9.2 Hz, colour coded as in Fig. 2E. **(C)** Amplitude spectrum for trials where a frequent target is preceded by an infrequent target. As in (A), the green line represents the mean vector length and the shaded area around ± 1 SEM. The mean number of trials in the *ifrq-frq* condition was 376 ± 25. **(D)** The individual vectors at 9.2 Hz, colour coded as in (B). **(E-F)** Amplitude spectrum and individual vectors at 9.2 Hz for trials where an infrequent target is preceded by a frequent target. The mean number of trials in the *frq-ifrq* condition was 378 ± 26. **(G)** *P*-values yielded by the 2D permutation test, not corrected for multiple comparisons. Again, the whole spectrum is shown for completeness, however, our one-back analysis focused specifically on the frequencies near the peak frequency, 9.1 Hz in Fig. 2C.

Finally, Figures 4E&F show the amplitude spectrum and individual vectors at 9.2 Hz for the *frq-ifrq* condition. Supporting our hypothesis, there was no significant oscillation in the *frq-ifrq* data (light-green dashed line in Fig. 4G). The amplitude at 9.2 Hz is 0.022 ± 0.014 and the vectors in Fig. 4F show a mean phase angle of 277° ± 7°. As the number of trials in *frq-ifrq* (Fig. 4E, *n* = 378 ± 26 trials) and *ifrq-frq* (Fig. 4C, *n* = 376 ± 25 trials) conditions were very similar (~20% of the trials), it is unlikely that the absence of significant oscillations in the *frq-ifrq* data was due to a lack of power.

The results of the one-back analysis align with our previous findings (Ho et al., 2019) that alpha oscillation (~9 Hz) of response bias is largely present on trials where the target occurs in the same ear as the previous target (*frq-frq*). Bias also oscillated at a similar frequency when a frequent target followed an infrequent target. Given that infrequent targets very rarely occurred on consecutive trials, the presence of a bias oscillation in the *ifrq-frq* data implies that it is likely linked to the frequent target two trials back. We verified this by pooling together the *ifrq-frq* and *frq-frq* trials. The GLM analysis revealed a clear oscillation in bias around 9 Hz with similar amplitude and phase angle as in the *ifrq-frq* and *frq-frq* data. Taken together the results suggests that the ear-specific alpha oscillation of bias lasts at least two trials, which is consistent with our earlier findings (Ho et al., 2019).

Why did we fail to observe an oscillation in the *frq-ifrq* data? As infrequent targets were rarely preceded by an infrequent target **two trials** back (i.e., *ifrq-frq-ifrq*), the *frq-ifrq* data contained largely trials on which the target switched ears, i.e., from 1-back *frequent* to current *infrequent* but rarely from 2-back *infrequent* back to the same ear in the current trial (see Fig. 1C).

### Rhythmic modulation of accuracy following an infrequent target

Although we failed to find any rhythmic modulation of accuracy when we collapsed across frequent and infrequent targets (Figs. 2A&B), there could be oscillatory effects associated specifically with infrequent (oddball) targets, which typically elicit an ERP difference wave, termed a Mismatch Negativity, associated with *prediction error* (Garrido et al., 2009; Stefanics et al., 2014; Wacongne et al., 2012; Winkler, 2007; Winkler et al., 2009). We applied the same one-back analysis to the accuracy data as we did to the response data. As shown in Figure 5C, there was a robust oscillation of accuracy around 7.6 Hz, *p* = 0.02 (corrected for multiple comparisons), in the *ifrq-frq* condition, with an amplitude of 0.017 ± 0.006. Neither the *frq-frq* (Fig. 5A) nor the *frq-ifrq* data (Fig. 5E) showed a similar peak in the amplitude spectrum. Correspondingly, no other significant frequency was found by the 2D permutation test shown in Figure 5G (same as in Fig. 2F). Figure 5D shows the vectors of individual subjects at 7.6 Hz in the *ifrq-frq* condition, which cluster coherently around a mean phase angle of 306° ± 12°. In comparison, the vectors at 7.6 Hz in the *frq-frq* (Fig. 5B) and *frq-ifrq* condition (Fig. 5F) are more uniformly distributed are more uniformly distributed, with vectors in all quadrants.

**Figure 5.**
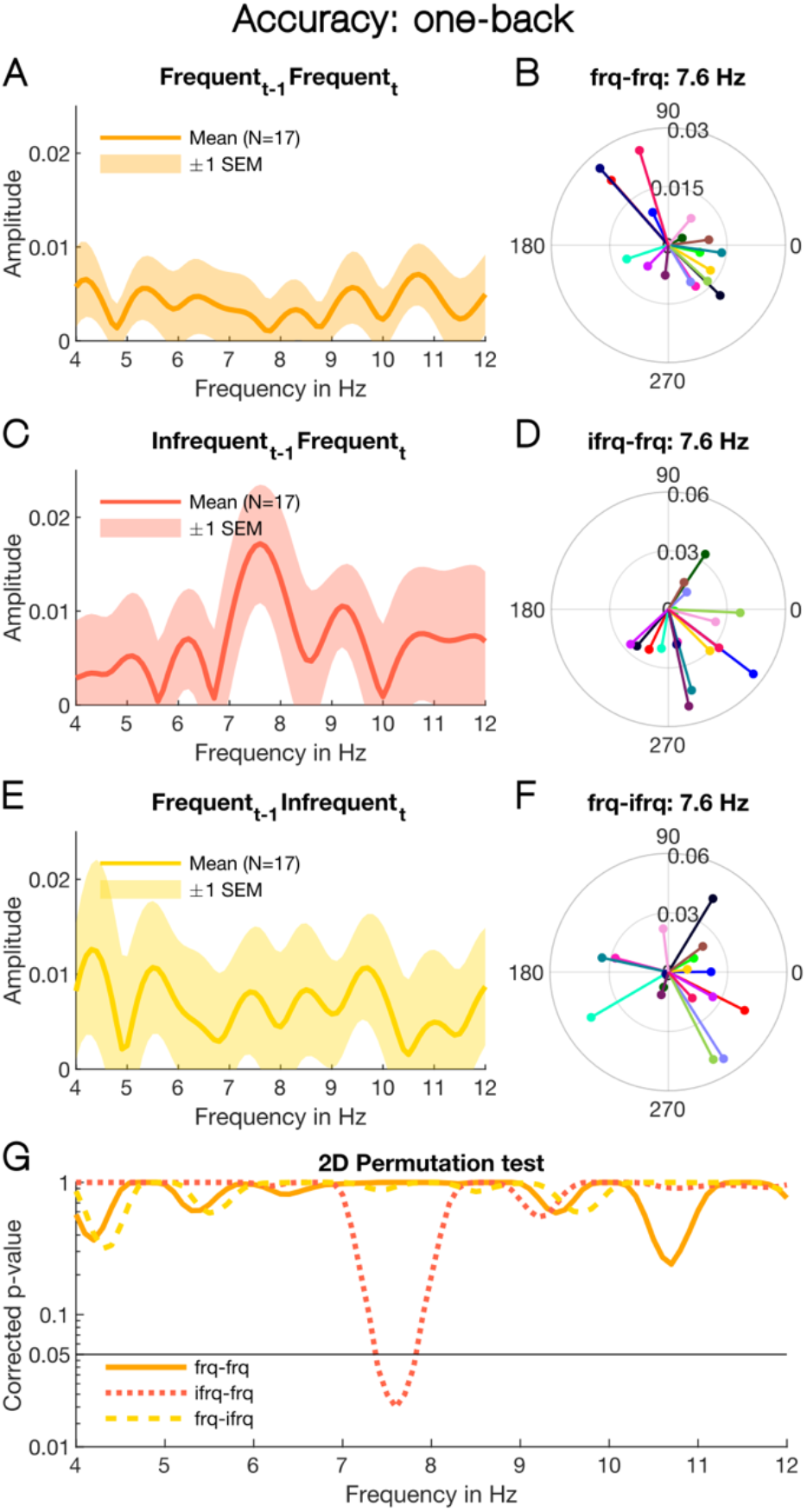
One-back analysis of accuracy. **(A)** Amplitude spectrum of accuracy for trials where frequent targets occur on consecutive trials: The orange line reflects the mean vector length across individual subjects at the tested frequencies and the shaded area indicates ± 1 SEM (N =17). **(B)** Vectors for individual subjects at 7.6 Hz, colour coded as in Fig. 2E. **(C)** Amplitude spectrum for trials where a frequent target is preceded by an infrequent target. As in (A), the red line represents the mean vector length and the shaded area ± 1 SEM. There is a clear peak in the spectrum around 7.6 Hz. **(D)** The individual vectors at 7.6 Hz, colour coded as in (B). **(E-F)** Amplitude spectrum and individual vectors at 7.6 Hz for trials where an infrequent target is preceded by a frequent target. **(G)** *P*-values yielded by the 2D permutation test, corrected for multiple comparisons using the same procedure as illustrated in Fig. 2F. The horizontal black line indicates *α* = 0.05. The mean number of trials in each condition was the same as in the analysis of response bias (Fig. 4).

The accuracy oscillation at 7.6 Hz shows several interesting characteristics. First, it appears to be specific to the oddball paradigm, as we have not encountered it in our previous study where we used the same task but presented the target with equal probability (50 %) to each ear (Ho et al., 2019). Moreover, we only found the oscillation when we separated the trials that were preceded an infrequent target from the trials preceded by a frequent target. This suggests that the oscillation in accuracy arises only after the occurrence of an infrequent target. Given the strong association of the oddball with prediction errors (Garrido et al., 2009; Stefanics et al., 2014; Wacongne et al., 2012; Winkler, 2007; Winkler et al., 2009), the 7-Hz oscillation we observe here could reflect the rhythmic behavioural modulation of predictive mechanisms over time.

## Discussion

### Propagation of auditory perceptual priors via alpha oscillations

Perception is a proactive process in which the brain uses past sensory information to generate predictions about the world. We have previously presented evidence that auditory perceptual history is propagated via a reverberatory mechanism oscillating at an alpha rhythm of 9-10 Hz (Ho et al., 2019). The present study extends our earlier findings by showing that this predictive oscillatory process functions in the same manner under conditions of high stimulus expectation. As in our previous study, participants were asked to identify the ear-of-origin of a monaural pure-tone target masked by uncorrelated, dichotic white noise. Stimulus expectation in each block was manipulated by increasing the target probability in one ear to 80 % of the trials. Decision bias was not constant across time but fluctuated rhythmically at ~9.1 Hz from noise onset, irrespective of whether the target occurred more frequently in the left (L80) or right ear (R80). Importantly, this cyclic modulation of bias at alpha rhythm was dependent on stimulus history, as in our earlier study (Ho et al., 2019). The bias oscillation was largely observed for successive frequent trials (*frq-frq*), in which the target occurred in the same ear in immediate succession. Although we define stimulus congruency in terms of ear, stimuli confined to one ear are perceived as originating from that side of space. Therefore, the interaction between consecutive stimuli may be mediated by the neuronal circuitry defining acoustic space, rather than within the monaural circuitry (King & Middlebrooks, 2011).

The ear or location specificity of the rhythmic modulation of bias suggests that no oscillation should be observed when the target switches ears. This was confirmed on trials where a frequent target was followed by an infrequent target (*frq*-*ifrq*). However, when an infrequent was followed by a frequent target (*ifrq-frq*), decision bias oscillated at ~8.8 Hz, only slightly slower than the 9.2 Hz observed for bias on the *frq-frq* trials. How do we explain that even though the target switched ears on the *ifrq-frq* trials, decision bias still oscillated? The *ifrq-frq* condition mostly contains two-back trials that are congruent (i.e., *frq-ifrq-frq*), as infrequent targets rarely occurred on successive trials. In our previous study(Ho et al., 2019), the oscillation of bias was still detectable on two-back congruent trials, suggesting that the rhythmic communication of perceptual priors can last at least two trials. Therefore, the oscillation at ~8.8 Hz likely reflects the rhythmic influence of stimulus information (of a frequent target) two trials back, although there are too few infrequent trials to test this hypothesis directly.

The present results for decision bias align completely with our earlier findings (Ho et al., 2019), despite the unequal target probability between the ears (80 % vs. 20 %) in the current study, which induced stronger stimulus expectation in one ear (within a block of trials). This suggests that the auditory system relies on past sensory information to maintain perceptual continuity even under conditions of high stimulus expectation. The ear or location specificity of the bias oscillations here and in our previous study (Ho et al., 2019) is consistent with findings showing that the influence of stimulus history is spatially tuned (Cicchini et al., 2017; Fischer & Whitney, 2014; St. John-Saaltink et al., 2016). Though still controversial (Fritsche et al., 2017), the conditions under which serial dependence arises imply that perceptual experience and sensory input interact on perceptual rather than decisional processes (Cicchini et al., 2017; Fornaciai & Park, 2018). In line with this view, our results suggest that the propagation of prior information occurs early within the sensory circuit (either the ear circuitry or that defining spatial location).

Several possible mechanisms may generate the oscillation of bias. Findings by VanRullen and MacDonald (2012) show that brief visual events elicit long-lasting reverberatory responses at alpha rhythm, termed “perceptual echoes”, which are involved in short-term memory processes (Chang et al., 2017; Huang et al., 2018). As in vision, auditory events could elicit similar memory traces that reverberate at alpha rhythm for at least two trials within the sensory circuit in which they were elicited (Fig. 1C). We assume that the phase of each trace is reset at the beginning of every trial by the noise bursts in both ears and that the oscillation in decision bias arises from the interaction between the incoming sensory information and the congruent prior. Alternatively, the binaural noise bursts at the start of each trial may trigger a loop of reverberating signals between top-down memory information and bottom-up sensory input, with the delay of the reverberation generating an excitatory/inhibitory modulation of the sensory response.

The working memory literature is divided on whether information can be stored in early sensory areas (Gayet et al., 2018; Scimeca et al., 2018; Xu, 2018). Pasternak and Greenlee (Pasternak & Greenlee, 2005) discuss evidence showing that the regions of the brain involved in sensory processing of a given perceptual attribute are modulated by working memory specific for that attribute. In match-to-sample tasks, the interactions between working memory and stimuli are highly sensory specific (Pasternak & Greenlee, 2005). In particular, Gottlieb et al. (1989) showed that neural response to a specific tone in the monkey auditory cortex was enhanced after presentation of a sample of matched frequency. They suggest that the sample changes the synaptic neural efficacy transiently, enhancing the response to the test. A similar mechanism could underlie the rhythmic modulation of decision bias. The previous tone alters the synaptic efficiency in specific circuits, so a target to the same ear (or location) is amplified whereas a target to the other ear is unaffected.

However, sustained spiking activity, by which information is thought to be maintained in working memory (Arnsten, 2013), are rare in sensory cortices and more common and robust in association areas of parietal, frontal and temporal lobes (Leavitt et al., 2017). Therefore, the function of sensory areas in working memory tasks may be to encode incoming information, while its storage and control take place at higher levels, in particular, the prefrontal and posterior parietal cortex (Xu, 2017). Oscillations could play a crucial role in supporting the information flow between these regions (Fries, 2005, 2015; Salinas & Sejnowski, 2001; Varela et al., 2001), with different rhythms serving separate functions at various levels of processing. Jia et al. (2017) recently showed that echo responses at ~10 Hz to visual objects presented concurrently in left and right hemifields were modulated by a low-frequency oscillation in the delta to theta range (2-4 Hz), causing cyclic inhibitions of alpha, depending on which hemifield or object the observer attended to. The alpha responses were clearly specific to the processing of sensory information at each target location, while the low-frequency modulation reflected a higher-order mechanism responsible for shifting attention between the two potential target locations.

Similar functional divisions between different oscillations could underlie sensory predictions. Recent intracranial findings by Sedley et al. (2015) suggest that alpha oscillations may encode the *precision* of auditory predictions, while faster rhythms (> 30 Hz) encode prediction errors. Below, we discuss additional results from the current study that suggest theta oscillations operate at a higher processing level, possibly across brain areas, to ensure that predictions remain up to date.

### Updating auditory perceptual priors via theta oscillations

The unequal distribution of targets between the two ears had no effect on the oscillation in decision bias. However, our manipulation of listeners’ expectation did cause a rhythmic modulation of accuracy, which has not been reported before. This oscillation was *not* observable when we pooled across the frequent and infrequent targets (Fig. 2), but once we separated the trials depending on whether the previous and current target was frequent or infrequent (same one-back analysis as applied to decision bias), the oscillation could be detected robustly at ~7.6 Hz. This rhythmic modulation of accuracy appears to be specific to infrequent targets and emerging only *after* the occurrence of an unexpected stimulus (on *ifrq-frq* trials) but not at the time of occurrence (*frq-ifrq*). Given these characteristics, it seems unlike any other oscillation of accuracy observed before (Fiebelkorn et al., 2013; Landau & Fries, 2012). Therefore, the 7.6-Hz modulation may reflect a process *un*related to the *rhythmic attentional sampling* mechanism that is thought to underlie oscillations of sensitivity, typically measured by accuracy (Fiebelkorn & Kastner, 2019; Landau, 2018; VanRullen, 2016).

Rhythmic fluctuations of sensitivity are often observed at 4-8 Hz (Benedetto et al., 2016; Fiebelkorn et al., 2013; Landau & Fries, 2012; Re et al., 2019; Tomassini et al., 2015, 2017) and are thought to arise from an attentional mechanism by which incoming sensory information is sampled in a cyclic manner, akin to a ‘blinking spotlight’ (VanRullen, 2016). Compelling EEG evidence from visual studies suggests that this rhythmic sampling mechanism involves ongoing brain oscillations in the theta frequency band (Busch et al., 2009; Busch & VanRullen, 2010; Mathewson et al., 2009), and several studies point to a similar oscillatory process in audition (Hickok et al., 2015; Ng et al., 2012). In a recent study, we showed that when participants had to discriminate the pitch of a brief monaural pure-tone target masked by uncorrelated, dichotic white noise, sensitivity in each ear oscillated at ~5-6 Hz but in *antiphase* (~180º). This is consistent with visual evidence of similar antiphase oscillations in accuracy between two potential target locations in left and right visual hemifield (Fiebelkorn et al., 2013; Landau & Fries, 2012).

The 7.6-Hz oscillation in accuracy that we observe in this study does not show this antiphase relationship between the two ears, otherwise the oscillations in left and right ear would have cancelled each other out. We showed this in our previous study (Ho et al., 2019) where we separated the trials depending on the target’s ear of origin and found that the vectors in the two conditions exhibited a tendency to pull in opposite directions. This explained the absence of accuracy oscillations when we pooled all trials together. Therefore, the 7.6-Hz accuracy oscillation we observe in the present study must have the same phase whether the target occurred in the left or right ear. This, and the fact that the oscillation emerged specifically after an infrequent target, dissociate it from the rhythmic sampling mechanism.

What process could the 7.6-Hz oscillation reflect instead? It may be related to the Mismatch Negativity (MMN), which is a brain response elicited by violations of perceptual regularity in the auditory (Näätänen et al., 2001, 2007) and visual modality (Stefanics et al., 2014). More specifically, the MMN is a negative deflection ~150-250 ms in the difference wave between the ERP to an infrequent target, also called a deviant, and that of a frequent target, or standard (Näätänen et al., 2001, 2007; Stefanics et al., 2014). Initially, it was considered an electrophysiological correlate of memory mismatch but is increasingly interpreted as a signal of *prediction error* that compels the perceptual system to update its predictions (Garrido et al., 2009; Wacongne et al., 2012; Winkler, 2007; Winkler et al., 2009). As updating predictions involves modifying sensory representations, this should affect perceptual sensitivity on subsequent trials (Alink et al., 2010; St. John-Saaltink et al., 2016; Summerfield et al., 2008; Todorovic et al., 2011; Wacongne et al., 2011). We propose that just as priors are propagated via alpha oscillations, their *update* could be communicated through theta rhythm, giving rise to cyclic modulations of accuracy following unexpected stimulus events.

Several studies show that MMN elicitation is associated with increases in theta power and phase coherence in frontal and temporal areas (Fuentemilla et al., 2008; Hsiao et al., 2009; Ko et al., 2012). Connectivity analyses suggest that fronto-temporal interactions and reciprocal changes are essential to MMN generation (Garrido et al., 2008), with theta oscillations playing a possible role in synchronising activity between frontal and temporal regions(Choi et al., 2013). The significance of theta rhythm in cortical coherence is corroborated by clinical findings, linking MMN deficits in Schizophrenia to decreased theta band activity (for a review, see Javitt et al., 2018). Schizophrenia is a well-known mental disorder that is characterised by episodes of psychosis. Theories of Schizophrenia range from dopamine and glutamate dysfunctions to structural and functional abnormalities (Insel, 2010). More recently, Bayesian approaches to Schizophrenia emphasise impairments related to probabilistic learning and prediction error and suggest that the hallucinations and delusions experienced by patients are due to deficient inference mechanisms that fail to integrate new evidence, resulting in false predictions (Fletcher & Frith, 2009; Griffin & Fletcher, 2017). Abnormal theta activity in Schizophrenia could contribute to this deficit.

Our finding is consistent with the idea that theta is involved in regulating the information flow between brain regions during the update of perceptual priors. Unlike the bias modulation, the accuracy oscillation we observe at ~7.6 Hz is *not* ear or location specific but results from a surprising switch of ears or locations. The MMN can also be elicited by a change in sound location, pointing to a higher-order change-detection process^7,8^. Taken together, the findings suggest that the accuracy oscillation observed here arises from the same mechanism (i.e., of detecting perceptual irregularities and updating sensory predictions) that underlies the generation of the MMN.

## Conclusion

We have identified two separate oscillatory mechanisms that operate concurrently to ensure auditory perception remains smooth and stable over time. They show that the auditory system engages in sensory prediction and relies on past perceptual experience to anticipate forthcoming sensory input, even under high stimulus expectation. The propagation of prior information involves alpha oscillations at ~9 Hz, which causes decision bias to fluctuate rhythmically. When a surprising event violates a prediction, the perceptual system updates its expectations via theta oscillations at ~7 Hz, which leads to rhythmic modulations of accuracy. Furthermore, alpha and theta rhythms operate at different levels of processing, with alpha modulating early processes at the sensory level, while theta oscillations mediate the information flow across sensory circuits at a higher level. In sum, oscillations play a crucial role in sensory predictions, with different rhythms serving distinct functions in the processing hierarchy.

## Acknowledgements

The research was supported by the Australian Research Council Discovery Project (DP150101731) and the European Research Council (H2020) 832813 “GenPercept” and MIUR PRIN 2017.

## Author Contributions

H.T.H. conducted the experiment, and M.C.M., D.C.B., D.A., and H.T.H. designed the experiment, defined and validated data analysis methods, and wrote the paper.

## Data Accessibility

The source data files for Figures 2–5 are available at https://doi.org/10.17605/OSF.IO/XWDN6.

